# Aesthetic Appreciation and Environmental Values: A Cross-Cultural Study of Teachers in 34 Countries

**DOI:** 10.64898/2026.08.07.743443

**Authors:** François Munoz, Jérémy Castéra, Franz Bogner, Pierre Clément

## Abstract

**Background:** The relationship between aesthetic appreciation and environmental values remains a critical yet under-researched area in environmental psychology. Although the Two-Major Environmental Values (2-MEV) model—encompassing preservation and utilization dimensions—serves as a standard framework for assessing environmental attitudes, the integration of aesthetic perception within this structure has largely remained unexplored. This study investigates the conceptual linkages across a diverse international sample to determine whether aesthetic appreciation functions independently of a traditional environmental value framework.

**Methods and Findings:** We conducted a large-scale, cross-sectional survey involving 11,800 pre- and in-service teachers across 34 countries. Participants’ environmental values were evaluated using the 2-MEV scale, while their aesthetic appreciation of nature and the built environment was assessed using Osgood’s semantic differential technique. Employing principal component analysis, hierarchical exploratory factor analysis (EFA), analysis of variance (ANOVA), and within-class analysis (WCA), we accounted for cross-national variations and evaluated response consistency.

The results demonstrate that aesthetic appreciation comprises two distinct dimensions— focusing separately on nature and the built environment—that operate independently of traditional preservation and utilization values, showing only weak correlations. Furthermore, while the overarching psychological structure remains consistent globally, our findings reveal significant cross-national variations in respondent scores, particularly concerning utilization-related values.

**Conclusions:** These findings establish that aesthetic appreciation constitutes a distinct psychological construct separate from conventional environmental value frameworks. The observed cross-cultural variability underscores the necessity of accounting for national and cultural contexts when designing environmental education programs. By leveraging a robust, comprehensive global dataset, this study provides a vital empirical foundation for integrating aesthetic and value-based dimensions into future environmental research and educational policy.

## Introduction

Aesthetic appreciation represents a broad spectrum of human perception including emotional aspects, such as terrifying or reassuring [1]. Aesthetics of nature can be rooted in religion (the Garden of Eden, where God offers nature) and in philosophy (Descartes claimed that science can “make us masters and owners of nature”). Extensive research has shown that aesthetic appreciation is a key component of an overall appreciation of nature [2–4]. Emotions and preferences determine the way people perceive, think and define their individual potential actions [5], which represent their environmental values [6,7]. The link between aesthetic appreciation and environmental values then concerns the way people can build environmental concern and citizenship. It is thus of primary importance to acknowledge this connection in a context of education (e.g., Gould et al., 2018; Nyberg, 2017; Stickney, 2020). We investigate here how aesthetic appreciation is related to environmental values among in- and pre-service teachers in an international, large-scale survey.

Naturalist philosophers have soon promoted an ecocentric appreciation of “nature” as devoid of human intervention, so that naturalist contemplation involves an observer who should not interfere with nature. The naturalist view grounds on a positive aesthetic experience [e.g. John Muir in 11]. The “environment” surrounds a reference point, typically a human being, so that it pertains to a more anthropocentric view concerning the use and management of natural resources. Because of the misuse and overuse of natural resources, the term “environment” more and more relates to negative aesthetic experience, especially leading to ecoanxiety and ecological grief [12]. However, whether and how aesthetic appreciation differs between nature and environment remain to be addressed.

Environmental psychology examines “the interplay between individuals and the built and natural environment” [7] and thus the nature of both anthropocentric and ecocentric views. The 2-MEV framework (Two Major Environmental Values model) is a widely used psychometric model in environmental psychology and environmental education [13–16]. It is designed to measure an individual’s environmental attitudes, values, and perceptions regarding the natural world. Unlike older models that treat environmental views as a single, linear spectrum, the 2-MEV framework evaluates how people balance the protection of nature against its human-centric use. Thus, the 2-MEV framework posits that independent utilisation and preservation dimensions underlie the variation of environmental values. Preservation values are relative to the protection of natural components from human alterations, and thus concerns a more ecocentric view of “nature” [17,18], also denoted as intrinsic values. Utilisation values are broadly defined relative to the use of environmental resources, and thus concerns a more anthropocentric view of “environment”, also denoted as instrumental values.

Recent research has further emphasized another concept of relational values [19]. These values refer to the meaningful connections, bonds, and responsibilities that humans share with nature. Unlike intrinsic value (nature has value simply for existing) or instrumental value (nature has value because it is useful to humans, like timber or economic resources), relational values focus on: (i) care, stewardship, and responsibility toward ecosystems, (ii) aesthetic, recreational, and spiritual ties to the natural world, and (iii) identity and place attachment (how nature shapes who we are and our sense of home or community). Aesthetic appreciation is thus considered as part of relational values [20]. In addition, it has been shown that relational values could represent a dimension independent from the anthropocentric-ecocentric divide [21,22]. However, there is no formal test to date of the linkage between aesthetic appreciation and the intrinsic-instrumental values as considered in the 2-MEV model. Based on a meta-analysis on the relational values of nature in empirical research conducted by Pratson, Adams, and Gould (2023), it is evident that diverse interpretations of the relational values concept persist among scholars. Further investigation in the field is essential before establishing a robust framework and instruments (Pratson, Adams, & Gould, 2023; Kleespies & Dierkes, 2020).

In terms of science teaching, aesthetic experiences are generating considerable interest. We investigate the issue in a context of education, as the aesthetic appreciation of teachers should impact the way the next generation appreciates nature and environment, and how they will act accordingly in future. Nyberg & Sanders [23] underlined the importance of aesthetic experience for learners in science in general (see also: Jakobson & Wickman, 2008; Magntorn, 2007) and more specifically in environmental education [26,27]. Those works are rooted into the Dewey’s point of view that science and aesthetic are clearly linked [28]. Aesthetic appreciation is now considered an important driver of learning experience in environmental education [29–31]. We thus posit that teachers play a central role in the pupils’ citizenship development, in terms of transferring environmental values, but understanding how teachers’ environmental values are connected to aesthetic appreciation has still not been addressed so far. We hypothesize that developing students’ scientific knowledge and engagement regarding pro-environmental attitudes [32] involves an aesthetic component [33].

To advance a research-based understanding of the complex links between human values and the natural world, this study investigates a central, yet under-explored, question in global environmental psychology: how is aesthetic appreciation related to environmental values in a cross-cultural context? To address this, we pursued two main research questions: First, do teachers’ appreciations of nature and the built environment consist of independent aesthetic components? Second, how do these components relate to the preservation and utilization values classically exposed in the 2-MEV model? We hypothesized that a positive aesthetic appreciation of nature significantly contributes to preservation preferences (Brady and Prior, 2020), while a more negative appreciation would correlate with utilization preferences, particularly in the context of global anthropogenic changes. To test these hypotheses, we conducted a robust quantitative analysis of responses from over 11,800 pre- and in-service teachers across 34 countries, using a comprehensive questionnaire that included items on both aesthetic appreciation and environmental values. This large-scale, international study provides a unique and powerful basis for exploring these critical relationships and their implications for sustainable development.

## Method

We designed a questionnaire investigating values and aesthetic appreciation related to nature and environment. The questionnaire was first built in the context of the European project Biohead-Citizen, and used afterward in other countries [34,35]. Here we submitted the questionnaire to 11,816 teachers and future teachers of 34 countries (table S1). We assigned each country to a standard code following the international ISO 3166 reference (from https://github.com/lukes/ISO-3166-Countries-with-Regional-Codes).

The respondents of the survey were all volunteers who gave informed consent to participate in the study. Our dataset complies with the definition of secondary use of existing data. Our data are de-identified, entailing that retrieving personal information from our dataset is impossible. The use of these data does not require approval from the university ethics committee. In addition, such committee did not exist at the time the Biohead-Citizen project was designed.

### Questionnaire items

#### Environmental values

Table 1 presents the items of the questionnaire measuring Preservation values, with prefix PRE, and Utilisation values, with prefix UT. Fifteen items were coded between “agree” and “disagree” [36], and two Osgood differential items opposed “to be used” and “to be preserved”, (denoted as UPENV_A70 and UPNAT_A78, regarding environment and nature, respectively). When needed, we reversed the item scoring so that they were correlated in the same direction with the Utilisation and Preservation dimensions [35]. We defined the items following Forissier [37], Clément [38] and Bogner & Wiseman [13,15], following the 2-Major Environmental Values (2-MEV) model. Repeated and independent studies have supported this framework [e.g., Milfont and Duckitt 39,for further information, see 21]. Our first English master version was translated into each national language. Parallel independent back-translations assured to reach a consensus and to limit misunderstanding. A test-retest protocol was applied to examine the consistency of answers with a delay of one month. Items were selected during a pilot phase to be as consistent as possible across countries. Currently the 2-MEV items are translated in many language versions all over the world, and thus provide a robust basis for assessment of environmental values and comparisons at an international scale.

**Table 1.**
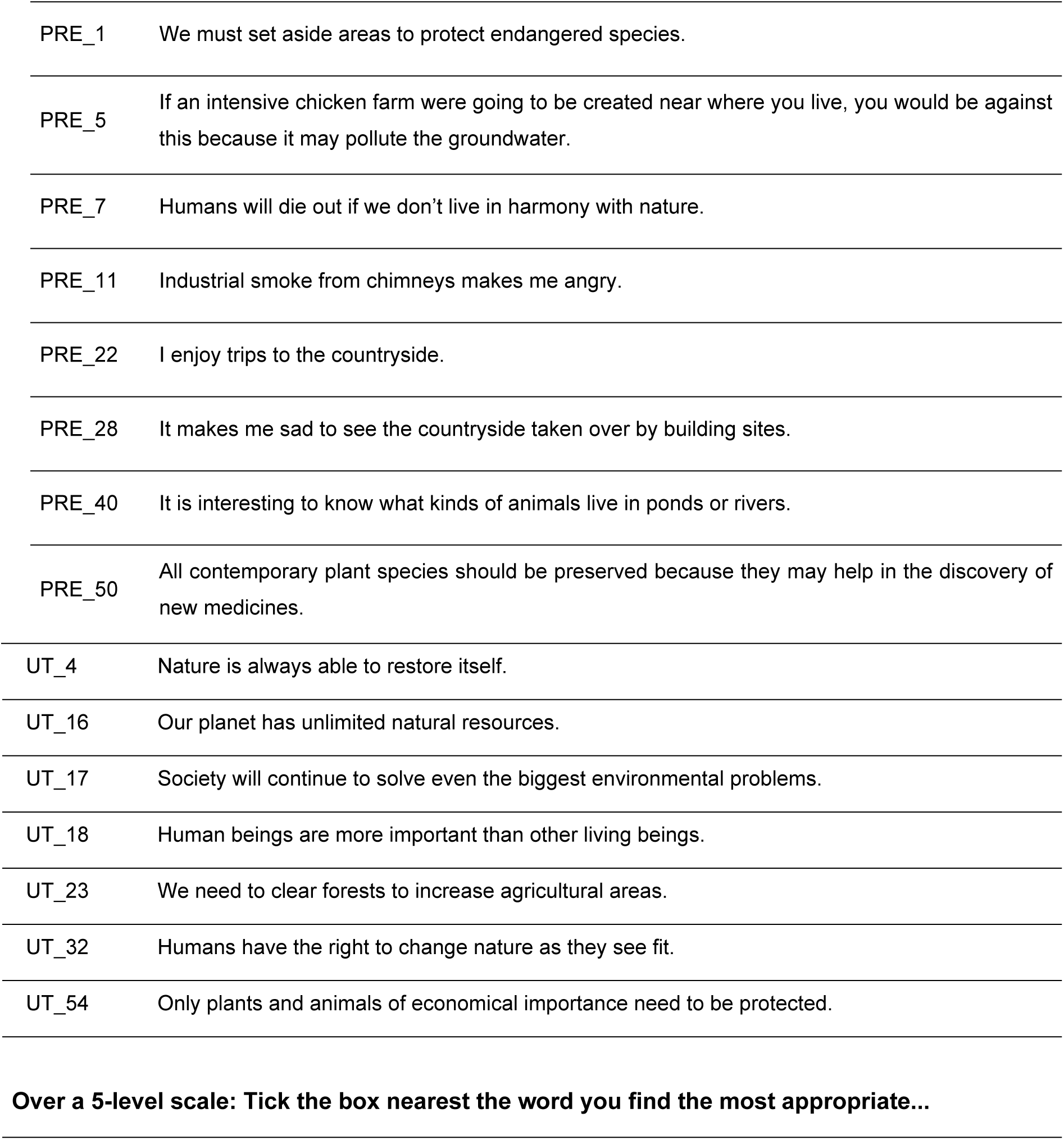
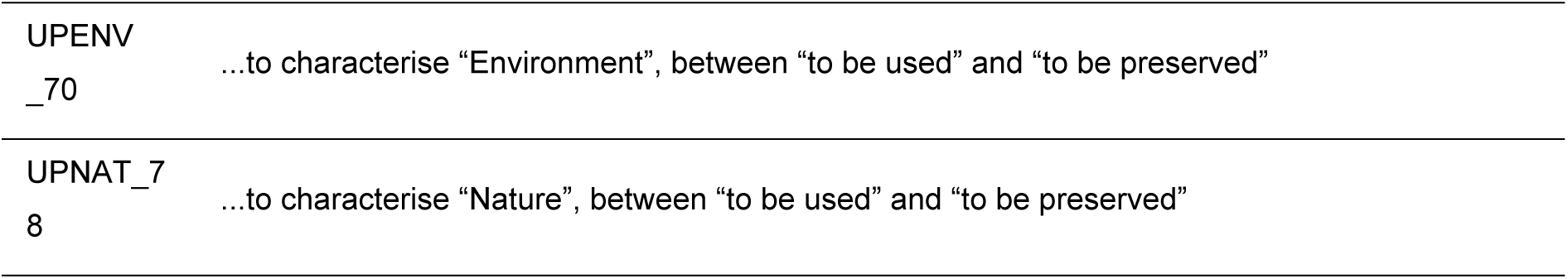
Environmental values items. . The items with PRE_ and UT_ prefix are related to Preservation and Utilisation dimensions of the framework, respectively. The answers are coded over a 4-level scale from “agree” to “disagree”. Items UTPR_A70 and UTPR_A78 request choosing a position closer to “to be used” or “to be preserved”, over a 5-level scale.

|  |  |
| --- | --- |
| PRE_1 | We must set aside areas to protect endangered species. |
| PRE_5 | If an intensive chicken farm were going to be created near where you live, you would be against this because it may pollute the groundwater. |
| PRE_7 | Humans will die out if we don't live in harmony with nature. |
| PRE_11 | Industrial smoke from chimneys makes me angry. |
| PRE_22 | I enjoy trips to the countryside. |
| PRE_28 | It makes me sad to see the countryside taken over by building sites. |
| PRE_40 | It is interesting to know what kinds of animals live in ponds or rivers. |
| PRE_50 | All contemporary plant species should be preserved because they may help in the discovery of new medicines. |
| UT_4 | Nature is always able to restore itself. |
| UT_16 | Our planet has unlimited natural resources. |
| UT_17 | Society will continue to solve even the biggest environmental problems. |
| UT_18 | Human beings are more important than other living beings. |
| UT_23 | We need to clear forests to increase agricultural areas. |
| UT_32 | Humans have the right to change nature as they see fit. |
| UT_54 | Only plants and animals of economical importance need to be protected. |
| **Over a 5-level scale: Tick the box nearest the word you find the most appropriate...** |  |
| UPENV<br>_70 | ...to characterise "Environment", between "to be used" and "to be preserved" |
| UPNAT_7<br>8 | ...to characterise "Nature", between "to be used" and "to be preserved" |

#### Aesthetic appreciation

To robustly investigate the relationship between aesthetic appreciation and environmental values, aesthetic appreciation is defined as a multi-faceted cognitive and affective process blending perceptual qualities, emotional responses, and evaluative judgments toward both natural and human-altered environments. Guided by this theoretical framework, we designed survey items to capture dimensions such as beauty, wonder, and sensory experiences, with content validity established through a clear link between construct definitions and questionnaire items. To assess these traits along a negative-to-positive continuum, we used the Osgood and colleagues’ Semantic Differentiation technique [40], employing a streamlined scale of 8 adjective pairs to effectively measure multidimensional affective aspects. The technique extracts individual choice between antagonistic terms representing affective aspects and defining the extremes of the scale (Forissier & Clément, 2003). Due to its simplicity, the method became popular in a variety of research disciplines and is regarded as an effective psychological tool to measure multidimensional responses [e.g., 42].

We considered a set of seven semantic differentiators with antonymous adjectives representing aesthetic appreciation of nature and environment. The items were derived from previous works of Clément et al. [43] and Forissier [37], following Quillot [44] for Nature and Larrère & Larrère [45] for Environment. Six items were related to aesthetic appreciation, ugly-beautiful (AENV_69 and ANAT_77), artificial-wild (AENV_ 71 and ANAT_79), unpleasant-pleasant (AENV_72 and ANAT_80), impure-pure (AENV_74 and ANAT_82), constructed-given (AENV_75 and ANAT_83), and bad-good (AENV_76 and ANAT_84). One was related to emotional dimension, terrifying-reassuring (AENV_73 and ANAT_81), which could evoke natural hazards such as earthquakes, volcanic eruptions, tsunamis, environmental pollution… but also some peaceful green landscape or beach. The same pairs were proposed to appreciate Nature and to appreciate the Environment. They were selected to be most discriminant among respondents [pilot test in France, Portugal and Germany, 37,41].

In the questionnaire version, we randomly set the order of the pairs of adjectives and, for each one, the order from negative to positive appreciation, to avoid biases in responses (Figure S1 in Supporting Information). For easier interpretation of the results, we subsequently recoded the responses to have consistent ranking from negative to positive appreciation.

### Sampling scheme

Respondents were secondary school pre- and in-service teachers of biology and national language, and pre- and in-service primary school teachers, thus forming six subsamples. In each country, each of the six groups included a minimum of 50 teachers, exceptionally fewer in some countries.

### Statistical analyses

Our statistical analysis was designed to rigorously address the relationships between aesthetic appreciation and environmental values on both a global and a cross-cultural level. We first analysed how the responses to aesthetic items for analogous pairs of words differed between nature and environment, by performing paired comparison tests for each item.

Next, we characterized the linkage of aesthetic appreciation items with values expressed by 2-MEV items. We performed a Principal Component Analysis of the global dataset including all items. We determined the number of relevant dimensions, *n*, by performing a scree-plot test [46]. Based on the number of relevant dimensions, we performed an Exploratory Factor Analysis (EFA) of the item structuring with a hierarchical design (Exploratory factor Analysis, Carroll, 1997). In this analysis, the global *g* factor defines any common structure of the items, i.e., a scale of appreciation/value expressed across all items. In addition, the analysis defines *n* orthogonal factors representing independent dimensions structuring the values expressed by respondents beyond the global common factor.

A key strength of this study is its multi-level analysis of cultural variations. We analyzed the variation across countries of the respondent scores on the factors of the EFA, by performing ANOVA analyses. In addition, we performed a within-class analysis of the item responses [wca function in R package ade4, 48]. This method is derived from PCA and allows characterizing the structure of responses within countries after accounting for the variation between countries. In order to check the consistency of the global structure of factors with the structure found within countries, we correlated the dimensions of the within-class analysis to the Principal Components performed on the whole dataset.

For all statistical analyses, we used R version 4.0.3 [49], with package *lavaan* for Factor Analysis [50] and *ade4* for within-class analysis [48].

## Results

### Aesthetic appreciation of Nature and Environment

Figure 1a shows the response patterns over the 5 levels of the 14 semantic differentiator items measuring aesthetic appreciation (as defined in Table 2). The items in the Figure are ordered by decreasing proportion of the most positive answer (greenest part). The 5 most positive items tend to appreciate nature as more beautiful (ANAT_A77), more pleasant (ANAT _80), better (ANAT_84), wilder (ANAT_79) and purer (ANAT_82). Conversely, the 5 items with least proportion in this category concern appreciation of environment, with greater relative importance of negative appreciation in orange, i.e., more constructed (AENV_ 5), more artificial (AENV_71), more impure (AENV_74), more terrifying (AENV_73), worse (AENV_76).

**Figure 1:**
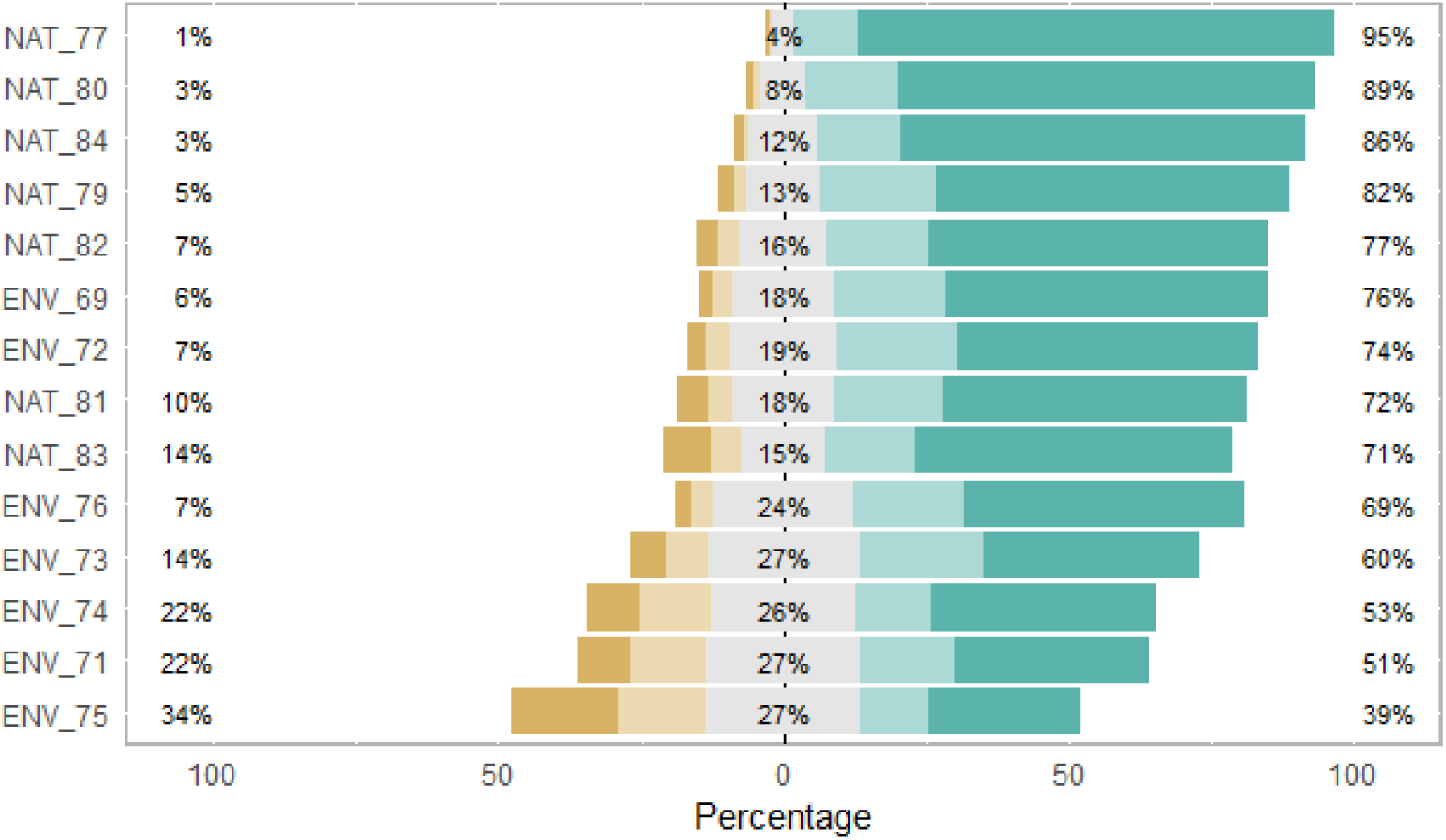
Aesthetic appreciation of Nature and Environment. The barplots include five coloured segments representing the percentages of responses to 5 levels of Osgood differentiator items. The items are shown, from top to bottom, by increasing proportion of answers in the median category. The item answers are coloured in green on the positive answer and orange in the negative part.

**Table 2.** Aesthetic appreciation items. . The same 7 pairs of adjectives are proposed to appreciate nature and environment, hence yielding 14 items.

| Over a 5-level scale: Tick the box nearest the word you find the most appropriate... |  |  |
| --- | --- | --- |
|  | ...to characterise<br>"Environment" | ...to characterise "Nature" |
| ...between "beautiful" and "ugly" | AENV_69 | ANAT_77 |
| ...between "wild" and "artificial" | AENV_71 | ANAT_79 |
| ...between "pleasant" and "unpleasant" | AENV_72 | ANAT_80 |
| ...between "terrifying" and "reassuring" | AENV_73 | ANAT_81 |
| ...between "pure" and "impure" | AENV_74 | ANAT_82 |
| ...between "constructed" and "given" | AENV_75 | ANAT_83 |
| ...between "good" and "bad" | AENV_76 | ANAT_84 |

We compared the structuring of analogous pairs of adjectives appreciating nature vs. environment. Figure 2 synthesizes the difference across respondents: the positive part represents respondents providing more positive appreciation of nature, while the negative part represents more positive appreciation of the environment. Many respondents (between 46 and 61%) provided the same response to analogous items. The remaining respondents tend to be more positive for nature than for environment (test of deviation from 0, all Student *p* < 0.001). Specifically, these respondents tended to find that nature is more beautiful, wilder, more pleasant, purer, more reassuring, more given and better than the environment.

**Figure 2:**
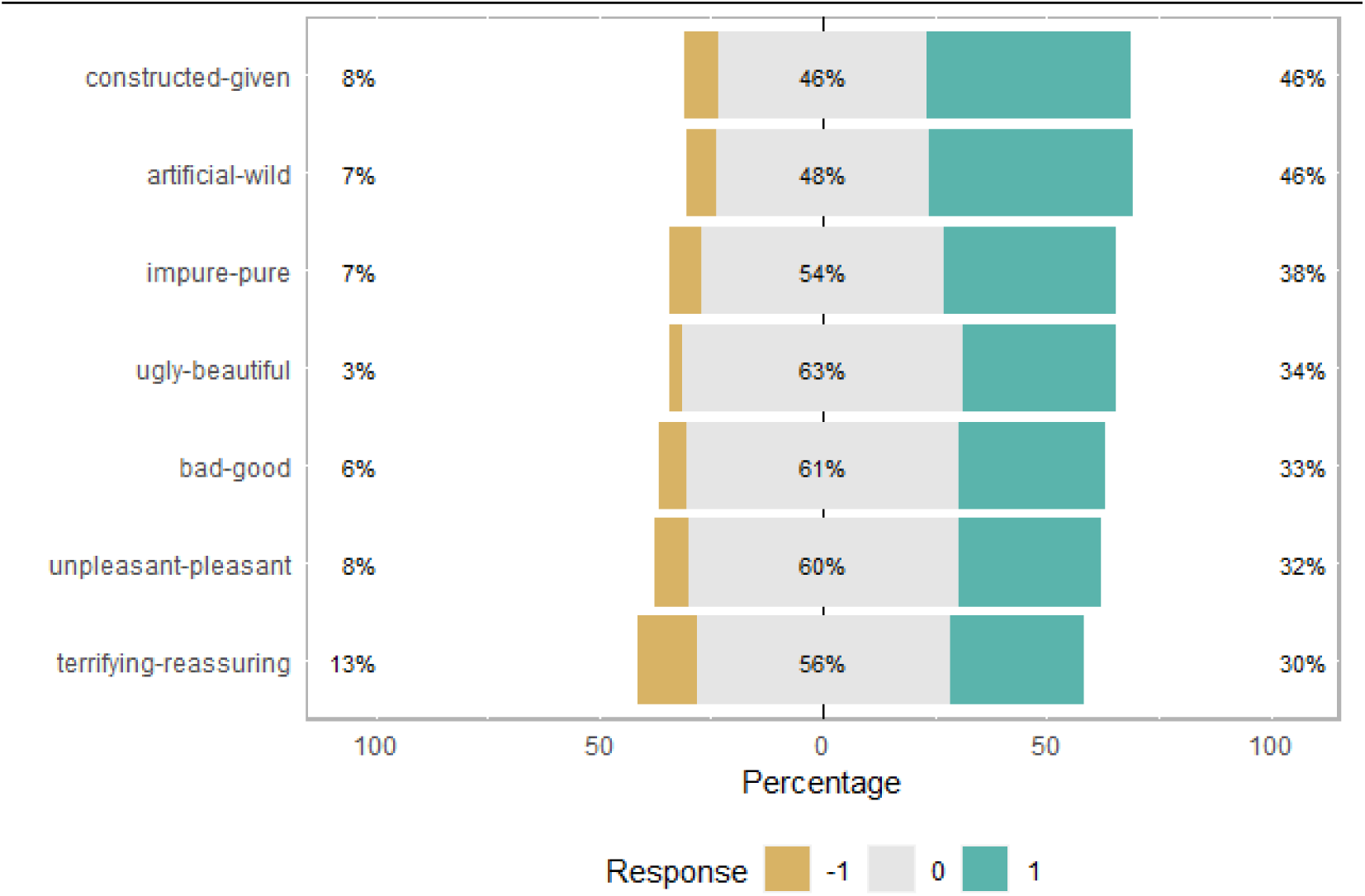
Difference between responses to analogous aesthetic items measured as the difference of nature score minus the environment score (orange if negative, green if positive). The class 1 thereby means more positive appreciation for Nature item, class −1 more negative answer to Environment item, and 0 means same answer to analogous Nature and Environment items.

### Relationship of aesthetic appreciation with environmental values

We found greater percentage of responses on the preservation side for both Osgood items scaling between to be used and to be preserved for Nature and Environment (Figure 3a). In addition, we found greater percentage on preservation side for Nature than for Environment (Figure 3b).

**Figure 3.**
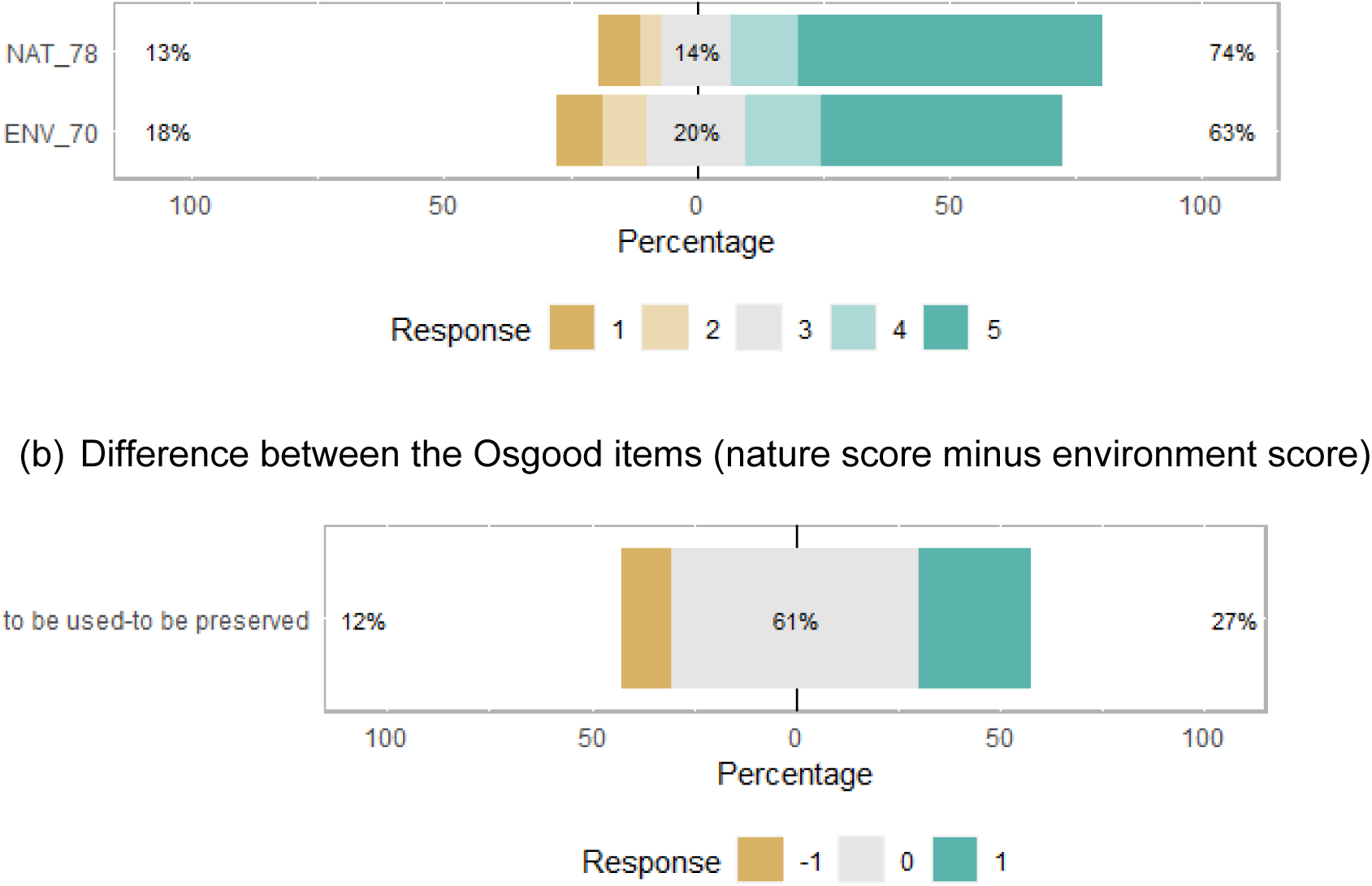
Osgood items contrasting Utilisation and Preservation values. The items scores from 1 (to be used) to 5 (to be preserved). (a) The barplots include five coloured segments representing the percentages of responses to the 5 levels. (b) Difference of nature score minus the environment score (negative in orange, positive in green). The class 1 thereby means more preservative value for Nature, class −1 means the reverse, and 0 means same answer for Nature and Environment.

We performed a Principal Component Analysis of all items, encompassing both aesthetic appreciation and environmental values. Based on the scree test, we identified four components with an eigenvalue neatly greater than 1 (i.e., above 1.6; subsequent components were below 1.15). Figure 4 then shows the hierarchical structuring of an Exploratory Factor Analysis (EFA) performed with four factors. On the left, the individual items can be related to each of the four independent latent factors F1 to F4. On the right, the global factor *g* expresses a global variation linking these latent factors [47]. The numbers on the arrows represent the correlation among the components across this hierarchy.

**Figure 4.**
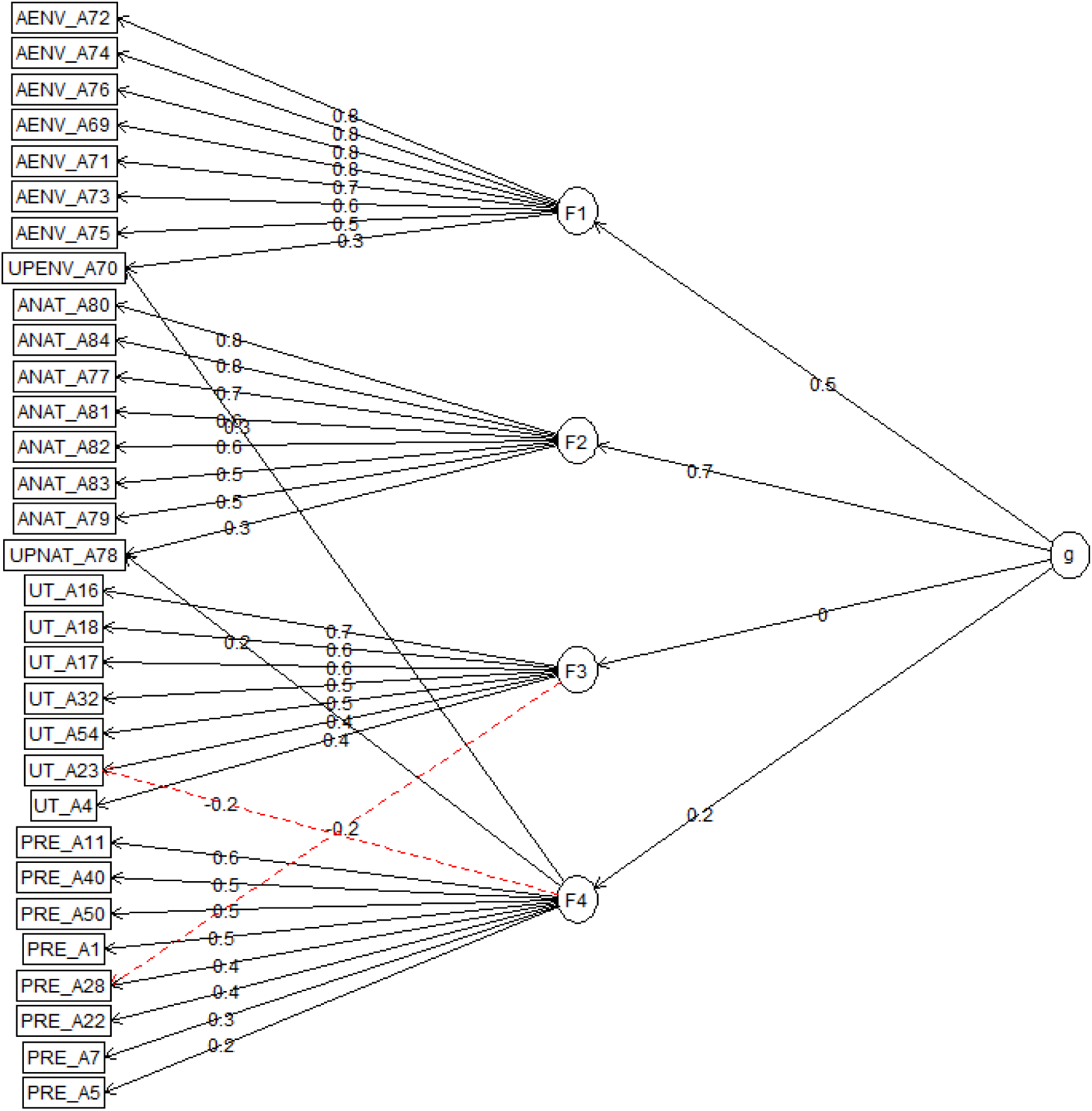
Second-order exploratory factor analysis for the sets of items concerning Preservation (PRE), Utilisation (UT) and emotional-aesthetic items (AENV and ANAT). The analysis yielded four basic orthogonal factors, F1, F2, F3 and F4. The values on the arrows represent the loadings of the items on the factors, with absolute value greater than 0.2 for individual items. The *g* factor represents a hierarchical structure above the four factors.

As also shown in Table 3, factors F1 and F2 were related to the Osgood items (mainly the aesthetic items), while F3 and F4 were related to items characterizing environmental values (both Likert and Osgood items). Furthermore, F1 related to environment items and F2 to nature items, and both scaled from positive to negative appreciation. The separation of environment and nature items on independent factors F1 and F2 likely results from the asymmetry of responses to analogous items (Figure 2), supporting the idea that aesthetic appreciations of nature and environment differ.

**Table 3.**
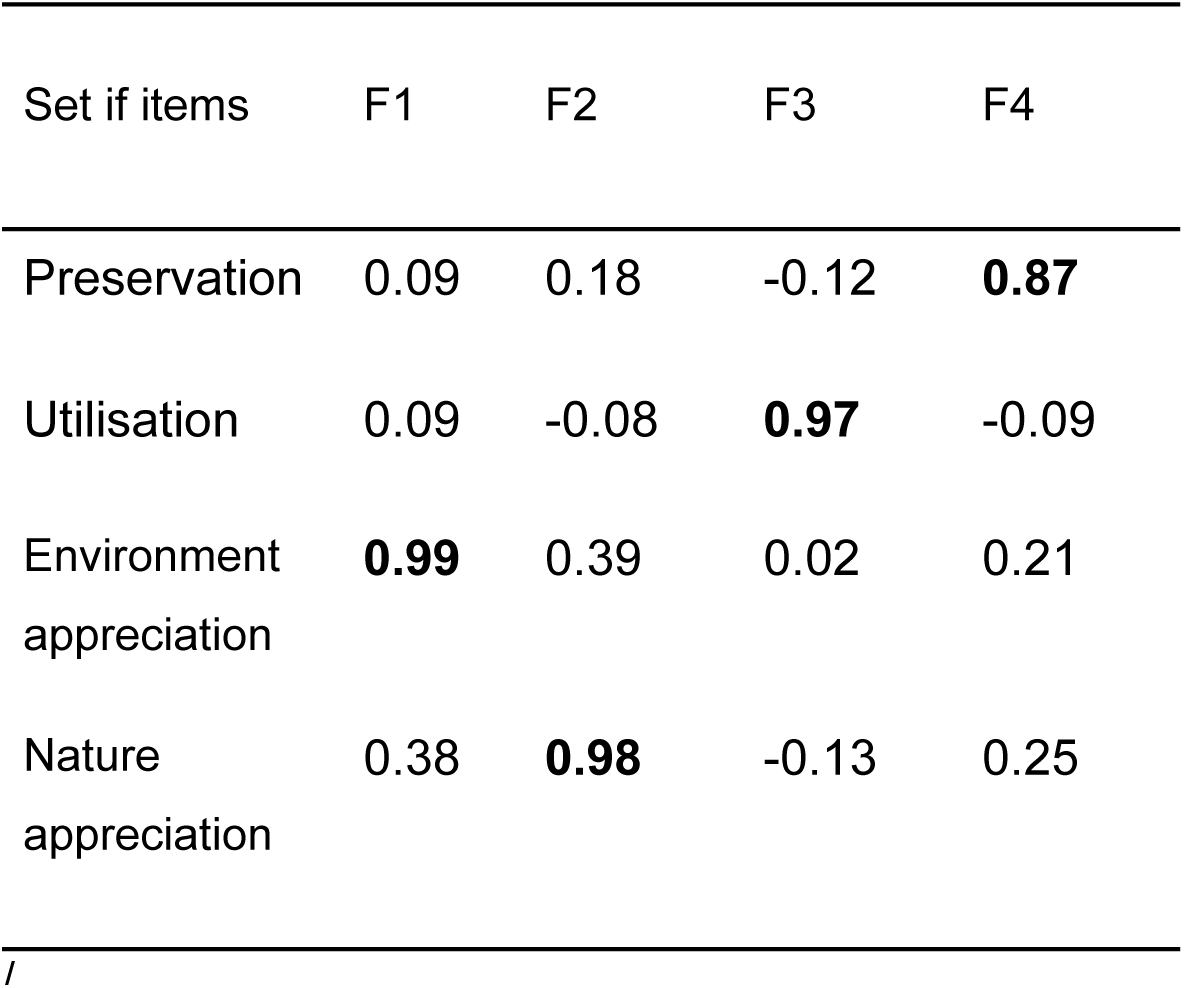
Correlations (Pearson) between the mean responses of teachers to four sets of items and the four factors (F1 to F4) defined from the Exploratory Factor Analysis. The highest correlation for each set of items is shown in bold.

F3 concerned utilisation items and F4 preservation items. The UPENV_A70 and UPNAT_A78 items scaling between to be used and to be preserved were both related to F1/F2 and to F4, indicating that they expressed conceptions related to both aesthetic appreciations and to preservation value.

The global factor *g* was primarily correlated to F1 and F2 but not to F3 and very little to F4: *g* underlined a common underlying structuring among all aesthetic items regarding nature and environment, and only a minor linkage between aesthetic and preservation items.

Although Utilisation and Preservation items globally formed two separate dimensions, UT_A23 (We need to clear forests to increase agricultural areas) was also related to the Preservation dimension, and PRE_A28 (It makes me sad to see the countryside taken over by building sites) was also related to the Utilisation dimension. These negative correlations indicate that pro-preservation values could be at least partly related to anti-utilisation values. Thus, partial opposition of utilisation and preservation values exists apart from the global decoupling of the dimensions F3 and F4.

### Structure of aesthetic appreciation and environmental values across countries

We performed ANOVA of the EFA score variation across countries (Figure S2 in Supporting Information). We found strongest variation for the F3 factor of utilisation values (R2 = 0.32, p < 0.001), followed by the F4 factor of preservation values (R2=0.14, p < 0.001), and the F1 factor of aesthetics of environment (R2 = 0.12, p < 0.001). The variation between countries was least marked but still significant for the F2 factor of aesthetics of nature (R2=0.05).

Furthermore, we performed a within-class analysis of the item responses to check the consistency of their structuring across countries [51]. The correlation between the first axis of within-class analysis and the first Principal Component of the global analysis was ρ = 0.999 (p < 0.001), between second axis and second Principal Component was ρ = 0.962 (p < 0.001), between third axis and third Principal Component was ρ = 0.877 (p < 0.001), and between fourth axis and fourth Principal Component was ρ = 0.551 (p < 0.001). This indicates that the PCA performed on all 34 countries captured a structure that is found in each country, independently from the variation between countries. Therefore, the structure of aesthetic appreciation and environmental values was broadly consistent across countries, suggesting a cross-cultural consistency of their dimensions.

## Discussion

Our study provides novel insights on the way aesthetic appreciation differs concerning nature and environment, and on the linkage between aesthetic appreciation and environmental values. Based on an unprecedented set of >10000 respondents from 34 countries, we identified several fundamental and independent dimensions, with a clear divide between preservation-utilisation environmental values and aesthetic appreciation (Figure 4). Furthermore, while many respondents provided identical aesthetic appreciation of nature and environment, many others still showed more positive aesthetic appreciation of nature (Figure 1). Although nature and environment are often considered equivalent in works on environmental aesthetics [e.g., 52], we demonstrate here more subtle differences. We primarily found a variation in utilisation values across countries, and the 4 dimensions found in EFA proved consistent within countries. Our results thus shed light on universal determinants of the variation in aesthetic appreciation and environmental values, beyond the variation between countries.

### Dimensions of 2-MEV environmental values

We primarily retrieved and confirmed the generality of the independent Utilisation and Preservation dimensions of environmental values, for a broad dataset spanning 34 countries (F3 and F4 factors in EFA analysis). The independence could reflect the distinction between a more materialist perspective, related to the use and management of natural resources, and a more idealist conception of nature, related to the preservation of the most significant natural jewels. However, we still found partial opposition between utilisation and preservation values, as reflected by the negative contribution of a Preservation item to the Utilisation dimension, and by the negative contribution of a Utilisation item to the Preservation dimension (Figure 4). For part of the respondents, the environmental concern expressed through the preservation values could thus be opposed to excessive or bad utilisation of natural resources [see also 34].

### Dimensions of aesthetic appreciation

People relationship with nature and environment should build on aesthetic appreciation [53–55]. Aesthetic appreciation is expected to be related to environmentalism and thereby to environmental values [56]. Aesthetic appreciation has indeed played a primary role in the development of conservation policies and actions, as underlined by Aldo Leopold in his famous essay [57,58]. Part of the tradition of natural aesthetics is picturesque and refers to a conception of “disinterestedness”. This tradition considers that nature has some intrinsic and dramatic beauty that transcends a materialist perspective. Conservation strategies following the picturesque perspective tend to preserve striking-eye and pristine ecosystems from human activities that altered other ecosystems [4,17,45,59]. Despite the expected relationship, we found that environmental values and aesthetic appreciation were broadly disconnected in EFA. Only a weak correlation of the factor *g* to the factor F4 could indicate that aesthetic appreciation is partly linked to preservation values. This could indicate that part of the respondents enjoy pristine and more preserved nature, that is, devoid of human intervention [60]. The otherwise independent dimensions of aesthetic appreciation could express a more idealist view, while 2-MEV environmental values could rather concern how human beings actually influence and act on ecosystems. Independence between 2-MEV items and nature/environment appreciation can also indicate that the 2-MEV framework is not designed to grasp aesthetic determinants of respondents’ attitudes [13,21].

### Does aesthetic appreciation differ between nature and environment?

We found that many respondents provided identical aesthetic appreciation for items concerning nature and environment (most differences at 0 in Figure 2). The global factor *g* of the EFA analysis expressed such a common appreciation, as it was correlated to both Environment and Nature appreciation items (right arrows in Figure 4). However, we also found that many respondents expressed different appreciation of nature and environment, as expressed by independent factors F1 and F2 in EFA. When respondents expressed different appreciation, we mostly found more positive appreciation of nature (Figure 2). Positive aesthetic appreciation of nature was mostly related to spiritual aspects (pure, pleasant, beautiful, Figure 1), while less positive appreciation was for items referring to human alteration (constructed-given, artificial-wild). Although Nature is sometimes considered as a component of our environment [61], the distinction in aesthetic appreciation here suggests some idealisation of Nature without human beings [see also 44,38], while human activities pervasively shape and somehow alter the environment.

### Variation across countries and global structure

Our sampling scheme in 34 countries spanned a broad range of cultural contexts. We found significant variation of the four factors across countries, yet stronger regarding utilisation values (R2 = 0.32; Supporting Information). The variation of environmental values across countries was already reported and could reflect influence of different socio-economical and cultural contexts [35]. Specifically, more utilitarian values are found in less-developed countries (Figure S1c in Supporting Information). Conversely, the variation was sensibly weaker for aesthetic appreciation (R2 < 0.12), indicating that most variation was found within countries. As Latour [62] noted, there is a “contradiction between a unifying but senseless nature, on the one hand, and, on the other, cultures packed with meaning”, and we would expect that the appreciation of nature depends on religion and culture [44]. As a consequence, different languages are expected to express relational values in different ways reflecting cultural variations [63]. Our questionnaire was translated in many languages and should be sensitive to such influence. These points support a variation in aesthetic appreciation across countries with contrasting cultural contexts, but the within-class analysis also showed that the same four factors could express the variation of values and aesthetic appreciation within the 34 countries. Therefore, the structure of environmental values and aesthetic appreciations, as making independent dimensions, proved consistent in a broad range of socio-cultural contexts.

### Influence of methodological choices

We acknowledge that the generality of the findings discussed above remain conditional to the design of the questionnaire:

i. We used a framework of adjective pairs with Osgood coding to assess aesthetic appreciation of respondents. There are some limitations and perspectives with this approach. First, a popular alternative option to assess aesthetic appreciation is based on images [3]. A sound perspective would be to compare the appreciation based on images and with Osgood items. However, using images implies specific choices of certain kinds of nature or environment images. In case of cross-cultural studies, the use of images has to be considered with precaution, for instance the choice of the image’s dominant colour can be differently appreciated according to the respondents’ countries [e.g., 64]. Second, we chose a limited set of adjective pairs to encompass different facets of aesthetic appreciation. Initially a broader set of 23 adjective pairs was designed and tested in three countries [37,41]. The 8 pairs of adjectives used in the present work were selected among the initial pairs during a pilot test, so as to reflect greatest variation of appreciation of nature and environment amond respondents. A perspective would be to design more in-depth analysis of appreciations by including some additional adjective pairs.
ii. We used two scales of responses to the items, either a 4-level Likert scale, or a 5-level Osgood scale. The Likert scale was suited to measure (dis-)agreement to statements regarding preservation/utilisation, hence it was used primarily for assessing the environmental values. Conversely, the Osgood scale was suited to represent a gradient of appreciation between aesthetically-connoted words, hence it was primarily used for the aesthetic items [40]. In addition, we included in the questionnaire two items that explicitly represented a gradient between “to be used” and “to be preserved” over an Osgood scale. We found that these items were related to the factor representing the Preservation dimension in the EFA (F4 in Figure 3). First, it indicated that the independent dimensions did not reflect a statistical artefact of Osgood vs. Likert items: two items with an Osgood scale were related to the dimensions of environmental values primarily coded on Likert scale. This finding is in line with previous works showing that using both Likert and Osgood items in a survey does not yield statistical bias related the kind of scale [65,66]. Second, it indicated that the responses between “to be used” and “to be preserved” terms primarily reflected preservation values (F4), and not utilisation values (F3). This results underline that the intentions of respondents tend to preservation values in some environmentalist perspective [67].
iii. The questionnaire analyzed here represents a limited set of items from a broader questionnaire, and further information could be grasped from other items [68]. For instance, Castéra et al. [34] analyzed the connection between 2-MEV items and items more specifically regarding the use of GMOs. In addition, we focused here on environmental values (V), and further research could assess the linkage of the values (V) with knowledge (K) and practices (P) [KVP model, 69]. Noticeably, other works suggest an interaction between aesthetic appreciation of nature and acquisition of scientific knowledge [1,4]. In a broader perspective, understanding the way emotions and aesthetics of nature and environment influence attitudes and practices is critical to improve the design and enforcement of management policies [e.g., 53–55]. Aesthetic preservationism would justify care of both nature and environment [70].

### Perspectives for science education

We can expect that the values and aesthetic appreciation together play a role in the process of environmental education and in building citizenship of students. Other works showed that using several educational aesthetic-based framework during training sessions can contribute to develop environmental ethics [9,23,71]. In this perspective, our results provide an important conceptual baseline to better understand and improve education practices, not only rooted in utilisation and preservation values, but also acknowledging aesthetic appreciation of nature and of the environment. Carr [72] argued that through the arts, environmental education should to bring closer “moral attitudes” and “aesthetic sensibility” to environment. Even if the relationships are not strong between values and aesthetic appreciation in our results, an important perspective for environmental education will be to build a stronger link contributing to a more integrated citizenship.

## Additional Information

**Table S1.**
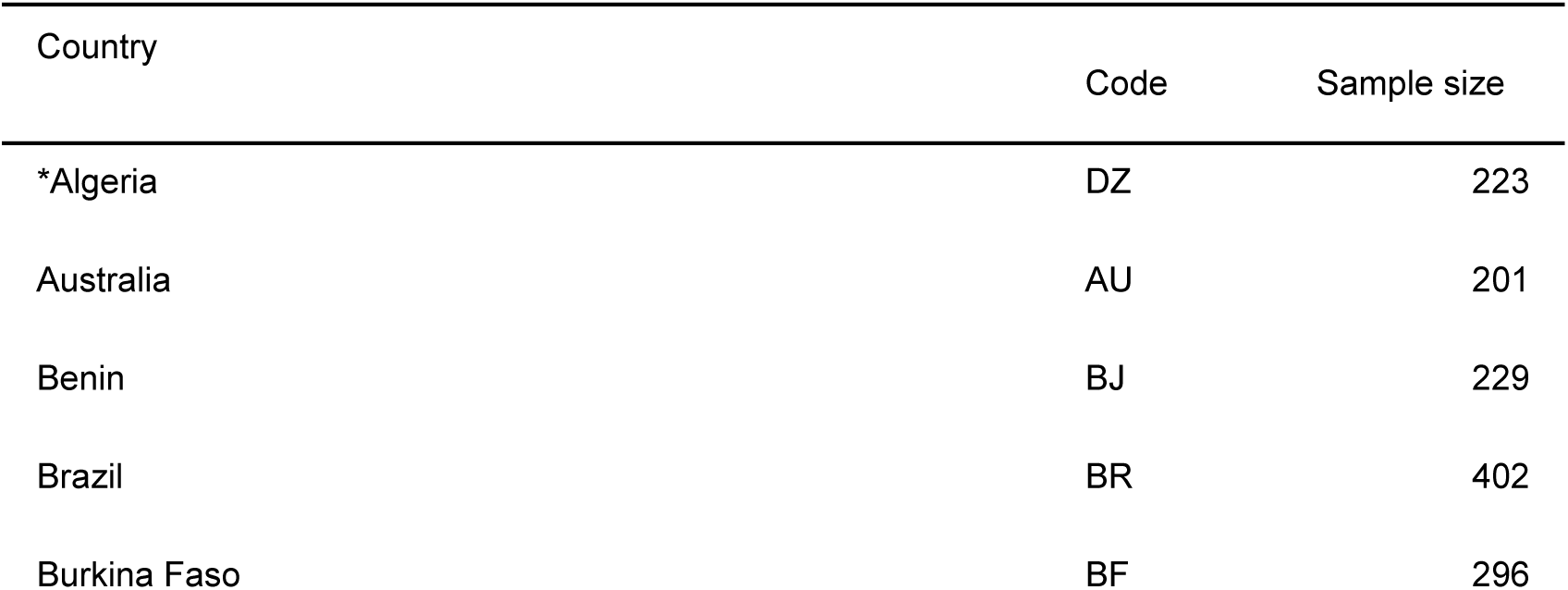
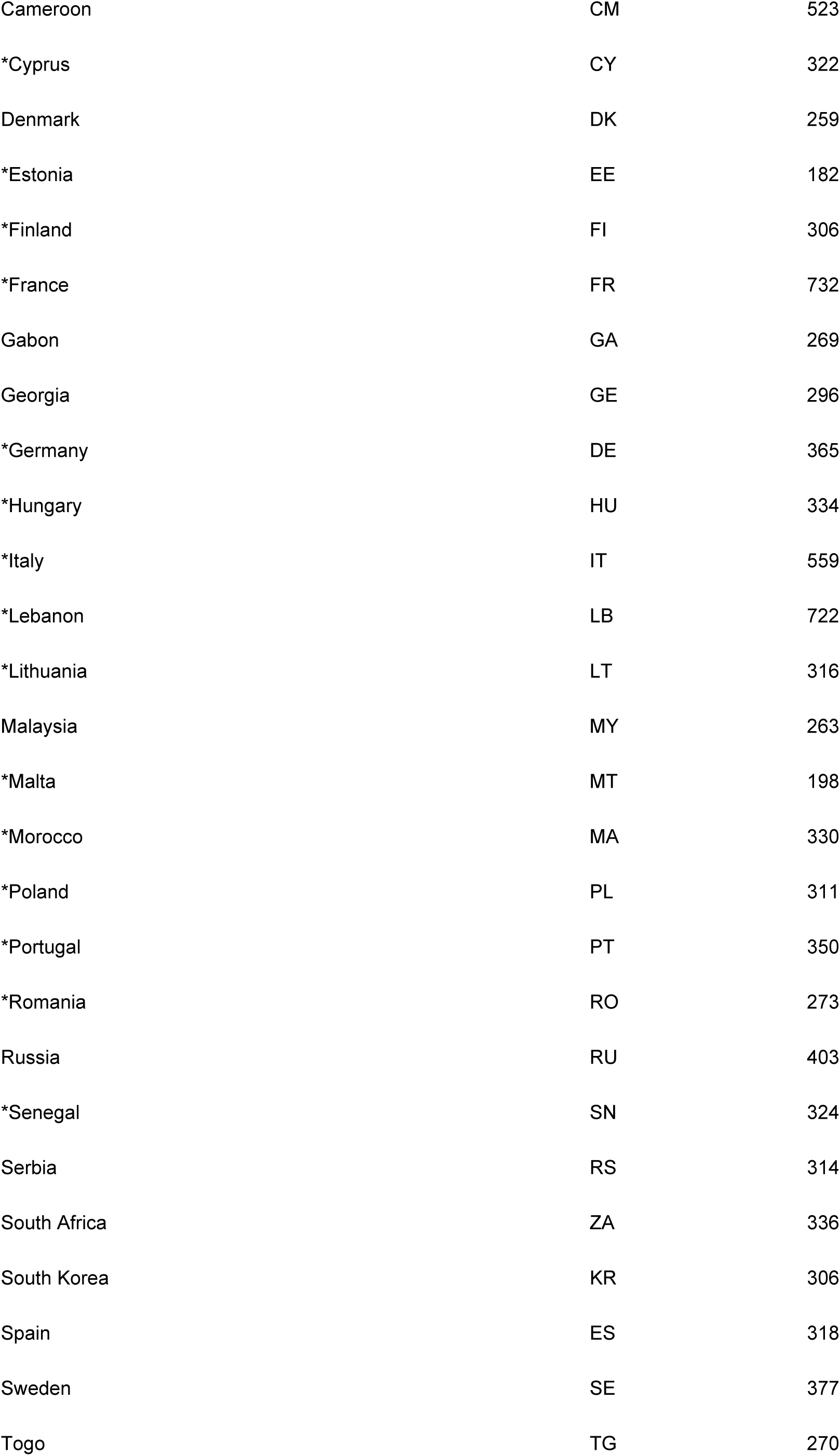
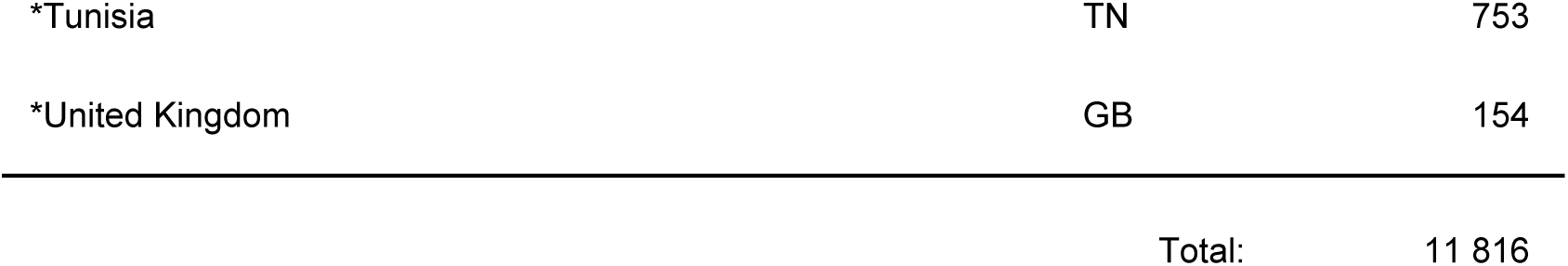
Number of respondents per country. * data 2006-2008, under the responsibility of Carvalho et al. [73] Other data (without *) were collected from 2009 to 2015 under the responsibility of P.Clément.

| Country | Code | Sample size |
| --- | --- | --- |
| *Algeria | DZ | 223 |
| Australia | AU | 201 |
| Benin | BJ | 229 |
| Brazil | BR | 402 |
| Burkina Faso | BF | 296 |
| Cameroon | CM | 523 |
| *Cyprus | CY | 322 |
| Denmark | DK | 259 |
| *Estonia | EE | 182 |
| *Finland | FI | 306 |
| *France | FR | 732 |
| Gabon | GA | 269 |
| Georgia | GE | 296 |
| *Germany | DE | 365 |
| *Hungary | HU | 334 |
| *Italy | IT | 559 |
| *Lebanon | LB | 722 |
| *Lithuania | LT | 316 |
| Malaysia | MY | 263 |
| *Malta | MT | 198 |
| *Morocco | MA | 330 |
| *Poland | PL | 311 |
| *Portugal | PT | 350 |
| *Romania | RO | 273 |
| Russia | RU | 403 |
| *Senegal | SN | 324 |
| Serbia | RS | 314 |
| South Africa | ZA | 336 |
| South Korea | KR | 306 |
| Spain | ES | 318 |
| Sweden | SE | 377 |
| Togo | TG | 270 |
| *Tunisia | TN | 753 |
| *United Kingdom | GB | 154 |

|  |  |
| --- | --- |
| Total: | 11 816 |

**Figure S1.**
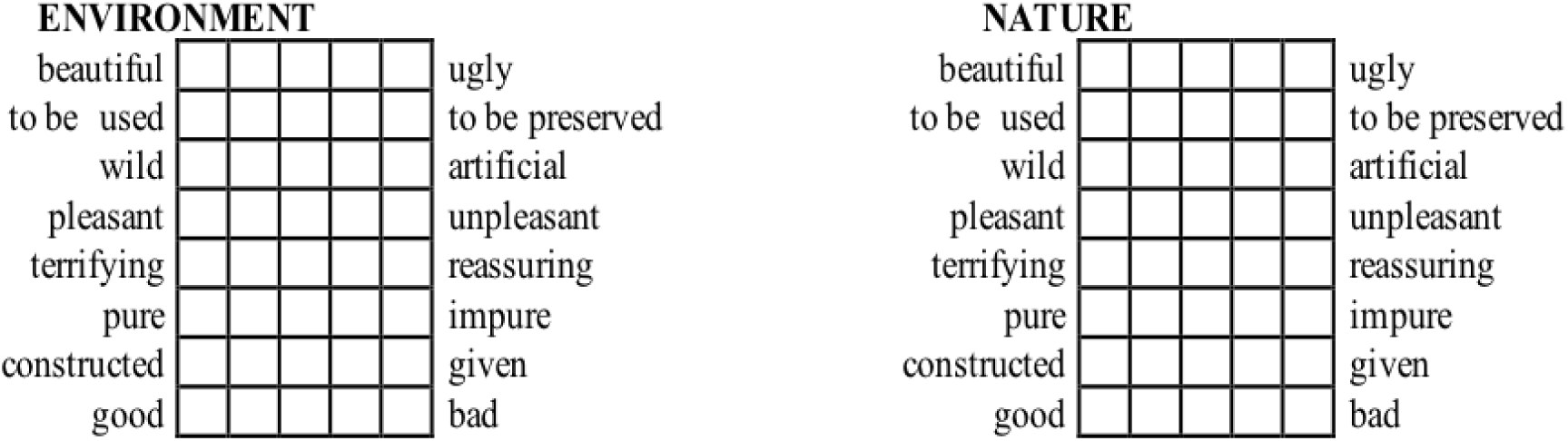
Original presentation of the Osgood-scales items in the questionnaire.

**Figure S2.**
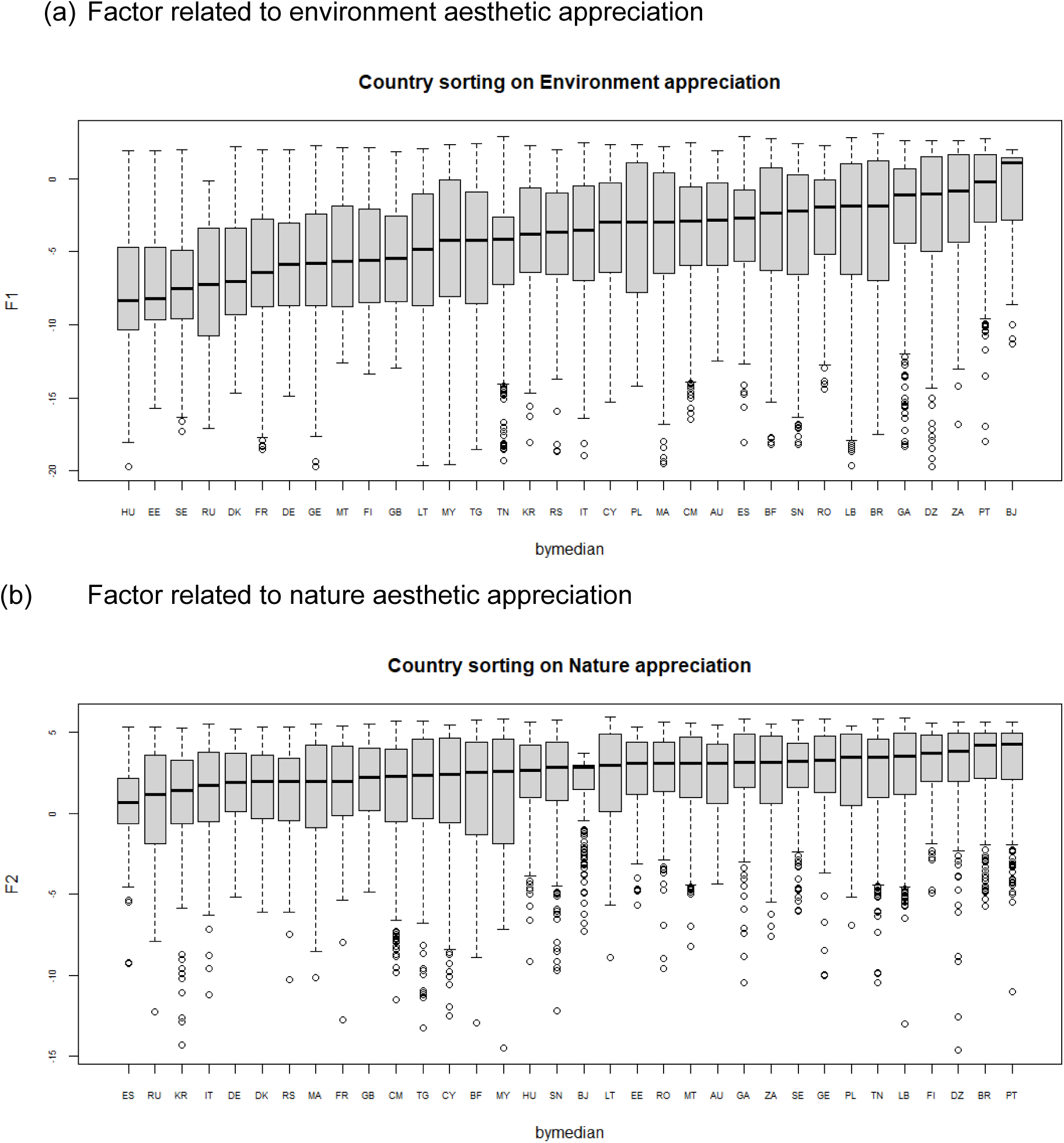
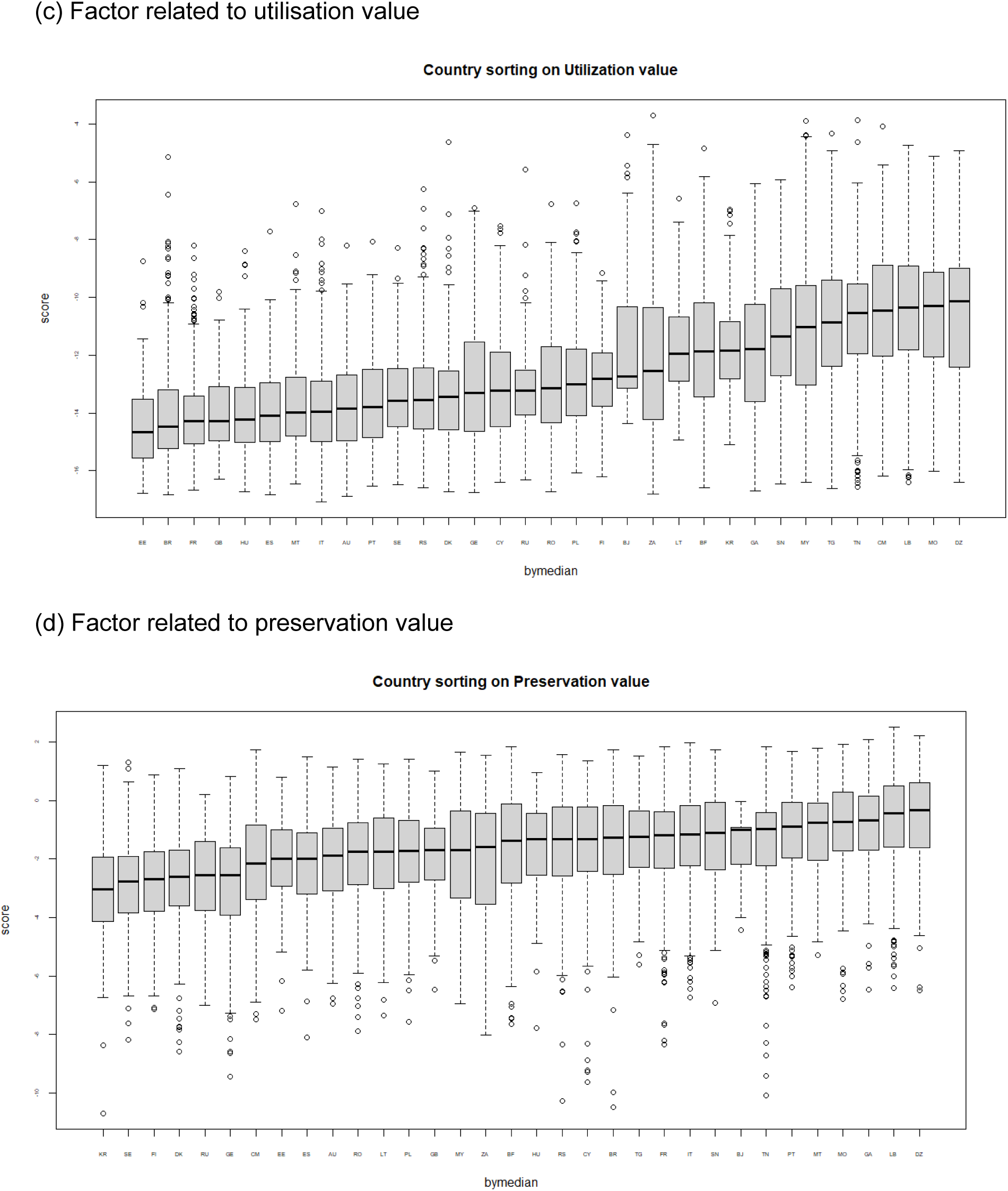
Boxplot of the four factors of the second-order Exploratory Factor Analysis (EFA), representing aesthetic/emotional components, (a) and (b), and environmental values, (c) and (d).

